# Elevated CO_2_ reduces a common soybean leaf endophyte

**DOI:** 10.1101/2021.04.06.438719

**Authors:** Natalie Christian, Baldemar Espino Basurto, Amber Toussaint, Xinyan Xu, Elizabeth A. Ainsworth, Posy E. Busby, Katy D. Heath

## Abstract

Free-air CO_2_ enrichment (FACE) experiments have elucidated how climate change affects plant physiology and production. However, we lack a predictive understanding of how climate change alters interactions between plants and endophytes, critical microbial mediators of plant physiology and ecology. We leveraged the SoyFACE facility to examine how elevated [CO_2_] affected soybean (*Glycine max)* leaf endophyte communities in the field. Endophyte community composition changed under elevated [CO_2_], including a decrease in the abundance of a common endophyte, *Methylobacterium* sp. Moreover, *Methylobacterium* abundance was negatively correlated with co-occurring fungal endophytes. We then assessed how *Methylobacterium* affected the growth of co-occurring endophytic fungi *in vitro*. *Methylobacterium* antagonized most co-occurring fungal endophytes *in vitro*, particularly when it was more established in culture before fungal introduction. Variation in fungal response to *Methylobacterium* within a single fungal operational taxonomic unit (OTU) was comparable to inter-OTU variation. Finally, fungi isolated from elevated vs. ambient [CO_2_] plots differed in colony growth and response to *Methylobacterium*, suggesting that increasing [CO_2_] may affect fungal traits and interactions within the microbiome. By combining *in situ* and *in vitro* studies, we show that elevated [CO_2_] decreases the abundance of a common bacterial endophyte that interacts strongly with co-occurring fungal endophytes. We suggest that endophyte responses to global climate change will have important but largely unexplored implications for both agricultural and natural systems.

## Introduction

Due to greenhouse gas emissions, the concentration of atmospheric CO_2_ ([CO_2_]) is currently ~410ppm and is forecasted to rise to 700ppm later this century – more than double pre-industrial levels and representing conditions that plants have not experienced for several million years (Leakey, 2009; Becklin *et al.*, 2017). Increasing [CO_2_] impacts plant phenotype in myriad ways, including development and phenology (e.g., flowering time) (Springer & Ward, 2007) and physiology (e.g., stomatal conductance, photosynthesis, transpiration) (Ainsworth *et al.*, 2002; Ainsworth & Long, 2005; Ainsworth & Long, 2020). The recent explosion in plant-microbiome work has emphasized that the microbes present in the leaves, stems, and roots of plants play critical roles in plant health and fitness (Trivedi *et al.*, 2020). Though plant physiological responses to CO_2_ have been the subject of hundreds of studies, the ways in which this and other aspects of a changing climate alter plant-associated microbial communities and thus plant health, both contemporarily and into the future, are just beginning to be unraveled.

Elevated [CO_2_] is known to affect ecological interactions between plants and other organisms, either by directly impacting plant mutualists or antagonists, or indirectly by modifying plant traits that mediate their interactions. For example, elevated [CO_2_] has been shown to increase pollinator visitation rate, potentially by modifying floral traits and volatile organic compound emissions (Glenny *et al.*, 2018). Plant-microbe symbioses in particular may be affected by increasing [CO_2_], as the functioning and fitness of the interacting partners are tightly linked (Becklin *et al.*, 2017). While on a geological timescale, periods of low [CO_2_] have created selective pressures for important adaptations in modern plants, there is little evidence for plant adaptation to contemporary increases in [CO_2_] (Leakey & Lau, 2012). However, because plant-associated microorganisms may adapt more quickly than plants to changes in climate (Lau & Lennon, 2012; Gehring *et al.*, 2017), shifts in microbial communities may be one way that plants can acclimate to elevated [CO_2_]. Our growing understanding of how plants respond to rising [CO_2_] has been facilitated by free-air CO_2_ enrichment (FACE) facilities, which manipulate greenhouse gas concentrations in the field (Ainsworth & Long, 2005; Ainsworth & Long, 2020). By combining data collected on plant-microbe interactions from these relevant field sites with controlled studies in the lab, we can improve our mechanistic understanding of plant-microbe and microbe-microbe interactions, and thus our ability to predict how these interactions will change as our climate does.

Elevated [CO_2_] can affect plant-microbe interactions in a variety of ways and through multiple mechanisms. For example, the majority of the studies on plant-mycorrhizae associations have shown that these interactions are positively affected by elevated [CO_2_] due to increased carbon availability to mycorrhizae, and increased plant nutrient demand (Alberton *et al.*, 2005, but see Kivlin *et al.*, 2013). Similarly, legume-rhizobium symbioses may also improve under elevated [CO_2_], as elevated [CO_2_] increases rhizobium abundance and symbiotic nitrogen fixation activity (Marilley *et al.*, 1999; Compant *et al.*, 2010), though this can be host specific (Schortemeyer *et al.*, 1996). Under elevated [CO_2_], rhizosphere community composition may also shift (Yu *et al*., 2016; Yu *et al*., 2018). For instance, fast-growing bacteria in the rhizosphere can take advantage of increased C rhizodeposition of plants to increase in abundance, as opposed to slow-growing bacteria that are not selected for under elevated [CO_2_] (Dorodnikov *et al.*, 2009). Most studies to date examining the effects of elevated [CO_2_] on plant-associated microbial communities have focused on belowground symbioses (summarized nicely in Grover *et al.*, 2015), despite the fact that leaf traits respond strongly to CO_2_ and may mediate adaptive evolutionary responses to elevated [CO_2_] (Leakey & Lau, 2012).

There have been only a few studies conducted on how microbes in aboveground tissues respond to elevated [CO_2_]. For example, the common fungal pathogen *Alternaria alternata* increases sporulation under elevated [CO_2_] (Wolf *et al.*, 2010). Under elevated [CO_2_] pathogenic *Colletotrichum gloeosporioides* has been shown to form more lesions (Pangga *et al.*, 2004) and in contrast, decrease in aggressiveness (the relative amount of damage caused to the plant without regard to resistance genes) (Chakraborty & Datta, 2002). Some cool-season grasses are colonized by systemic heritable fungal mutualists, and several studies have examined the impact of elevated [CO_2_] on those specialized aboveground symbionts and their grass hosts (Marks & Clay, 1990; Newman *et al.*, 2003; Ryan *et al.*, 2014; Chen *et al.*, 2017). However, aboveground plant tissues are also colonized more generally by diverse communities of symbiotic organisms. Changes to these microbial communities could have important consequences for plant acclimation to climate change, but such changes and their impacts are not well-understood.

All plant species are colonized by foliar endophytes: environmentally-acquired, asymptomatic leaf fungi and bacteria that can have important effects on plant physiology and ecology (Ryan *et al.*, 2008; Rodriguez *et al.*, 2009). For example, endophytes can affect plant nutrient acquisition and metabolism (Mejía *et al.*, 2014; Christian *et al.*, 2019), and offer protection from plant pathogens and herbivores (Arnold *et al.*, 2003; Estrada *et al.*, 2013, Busby *et al*., 2016). While the effects of individual endophytes on their hosts can be determined by inoculation experiments, recent research has also demonstrated that changes in overall endophyte community composition can have marked effects on plant health (Christian *et al.*, 2017). Endophyte community composition can be influenced by both biotic and abiotic factors, such as leaf traits (Van Bael *et al.*, 2017), location and spore dispersal (Christian *et al.*, 2016, 2017), and climate (U’Ren *et al.*, 2019; Oita *et al*., 2021). Because of this, there is growing interest in how endophyte communities will respond to climate change (U’Ren *et al.*, 2019; Rudgers *et al.* 2020), potentially in ways that can increase plant performance in the face of stressors such as drought (Giauque *et al.*, 2019) and UV-B radiation (Ramos *et al.*, 2018), yet there has been no such investigation of the impact of rising [CO_2_] on endophyte communities in field-grown plants. A recent study examined how elevated temperature and [CO_2_] interacted to affect endophyte colonization in highly controlled open-top chambers within a greenhouse, and found minor differences between chambers (Gonçalves *et al*., 2020). More studies, particularly those under natural field conditions, are needed to more fully understand how major drivers of climate change, such as elevated [CO_2_], impact how endophytes interact with their host plants and with each other.

Interactions among endophytes can shape microbial community composition and function, and play an important role in the health of the plant host. For example, order of arrival of a fungal endophyte and a fungal pathogen has been shown to affect the outcome of that interaction in lima bean (Adame-Álvarez *et al.*, 2014). Priority effects have also been shown to influence how synthetic communities of fungi assemble in *Populus* leaves in ways that affect host disease susceptibility (Leopold & Busby, 2020). Additionally, in a field manipulation experiment, priority colonization of cacao leaves by a dominant endophyte reduced overall diversity and density of the fungal endophyte community, in addition to reducing foliar pathogen damage (Christian *et al.*, 2017). This study suggested that priority colonization by the dominant endophyte induced expression of plant defense pathways that inhibited colonization or proliferation of microbes in general, including both endophytes and pathogens (Christian *et al.*, 2017). However, common endophytes could also have more nuanced interactions with the rest of the microbiome, acting as keystone species that either inhibit or facilitate other colonizers in ways that shape the distribution of species, or even strains, of endophytes within a leaf. *In vitro* assays have been used to examine how endophytes inhibit or facilitate pathogen growth rate, and similar methods could be useful for untangling how common members of the microbiome interact with other colonizers. Approaches that isolate competitive or facilitative effects in the lab would be especially powerful if combined with field surveys, allowing for organismal biology to be connected to patterns of community composition in the field.

Here we combine a field comparison of endophyte communities isolated from soybean (*Glycine max)* grown under elevated and ambient [CO_2_] with *in vitro* assays that test for the outcome of microbial interactions predicted from the field data. First, we capitalize on the Soybean Free Air Concentration Enrichment (SoyFACE) facility to test the effects of elevated [CO_2_] on endophyte communities inhabiting field-grown soybean. Because photosynthesized plant carbon is a resource for endophytes (Hardoim *et al.*, 2015), we predicted that endophyte abundance would be higher in plants exposed to elevated [CO_2_], due to elevated rates of host photosynthesis. Additionally, we predicted that endophyte diversity would be lower under elevated [CO_2_], due to selection favoring fast-growing endophytes that may outcompete slow-growing endophytes under elevated [CO_2_]. Finally, we predicted that due to direct effects of CO_2_ on endophytes, or indirect effects of CO_2_ on soybean growth or physiology, endophyte community composition would differ between plants grown under elevated [CO_2_] and ambient [CO_2_], despite being exposed to a similar endophyte species pool.

We then conducted three experiments to examine how *Methylobacterium* sp., the most abundant endophyte isolated from soybean in this study, interacted with soybean-associated endophytic fungi *in vitro*. Because *Methylobacterium* is known to be antagonistic towards fungal pathogens (Dourado *et al.*, 2015), we expected it to exert primarily negative effects on co-occurring non-pathogenic fungi. We also tested for the impact of establishment time by *Methylobacterium* on the outcome of the interactions, and expected that when *Methylobacterium* was more established, there would be more extreme effects on co-occurring fungi in culture. Finally, because surveys of community composition often focus on species or even genus-level shifts and might, therefore, miss functionally-important within-group variation, we tested for intra-operational taxonomic unit variation among fungal isolates in their response to *Methylobacterium*.

## Materials and Methods

### Field Site and Sample Collection

Field work for this study was conducted in June 2018 at the Soybean Free Air Concentration Enrichment (SoyFACE) facility located at the University of Illinois (40°02′N, 88°14′W; https://soyface.illinois.edu). Meteorological data for the 2018 growing season (May to October) can be found in Table S1. A commercial variety of soybean (*Glycine max* L.), Pioneer P32T16R, was planted on 18 May 2018 in 38.1 cm rows and plant density of ~52,500 plants per hectare. The field was treated with a pre-emergent herbicide (Authority Assist) and two post-emergent herbicides (Flexstar GT and AMS at recommended rates), consistent with standard agronomic practice in the region. Soybeans were grown in rotation with maize fertilized at ~200 kg ha^−1^; accordingly, no nitrogen was applied to the soybean crop in 2018. The field was not irrigated.

Four blocks, each containing two 20 m diameter octagonal experimental plots were established within a 16 ha soybean field. Each block contained one elevated [CO_2_] plot (600 μmol mol^−1^), and one control plot with ambient [CO_2_]. [CO_2_] in the elevated plots was controlled by an adjustable segmented octagon of pipes (20 m diameter) that released CO_2_ into the wind. Wind speed, direction, and CO_2_ concentration were automatically and continually recorded from the center of the plot, allowing the rate and position of gas release to be maintained at an elevated concentration within the plot (Rogers *et al.*, 2004; Eastburn *et al.*, 2010).

To investigate the effect of elevated [CO_2_] on endophyte communities, we collected leaves from 15 soybean individuals within each elevated [CO_2_] and ambient [CO_2_] plot from three of the four blocks. All plants were in reproductive stage R1 (beginning bloom). We collected one leaf from each plant (total n = 90 leaves). Specifically, we collected the youngest, fully expanded leaf from each plant individual. Relative age of leaves in elevated vs. ambient [CO_2_] plots was presumed to be similar. Leaves were free of visible pathogen and herbivore damage. Leaves were kept at 4°C and processed within 72 hours of harvest. At the time of sample collection, we also measured plant height.

### Endophyte Isolation and Identification

We used a custom 3D-printed stencil (design accessible here: https://www.tinkercad.com/things/llC8LHhTSYO) to excise two 0.8cm × 0.8cm fragments from the middle of each leaf, adjacent to the midvein. We dried one of these fragments at 65°C for three days to measure leaf mass per area (LMA). Following Christian *et al.,* 2016, we further subdivided the other fragment from each leaf into 16 pieces, each with an area of 4mm^2^. We surface sterilized the 4mm^2^ fragments as follows: Tissue fragments were submerged and agitated in 70% ethanol for 3 minutes, 0.0525% sodium hypochlorite for 2 minutes, and then sterile water for 1 minute. We then placed each piece of tissue in an individual Eppendorf tube containing 2% malt extract agar (MEA) (i.e., slant), media that selects for the growth of fungi. We chose MEA because we were originally focused on the isolation of fungal endophytes for this experiment.

We sealed slants with Parafilm and incubated them with a 12/12 h light/dark cycle at room temperature. Slants were incubated for over a year, but after 7 months no new fungi emerged from plant tissue. We subcultured emergent hyphae onto 2% MEA plates and allowed hyphae to grow until the colony covered the agar plate. We grouped subcultures into colony morphotypes based on color and mycelial growth patterns. With the exception of the most abundant fungal morphotype, we sequenced all fungal isolates. For the most abundant fungal morphotype, we selected 15 representative isolates for sequencing. We suspended vouchers of living mycelia for all isolates in sterile water and stored them at room temperature at the University of Illinois.

We extracted total genomic DNA directly from fungal isolates using a MO BIO (now Qiagen) PowerPlant® Pro DNA Isolation Kit. We used primers ITS5 and ITS4 to amplify the internal transcribed spacer (ITS) region of fungal DNA. Each 20 μL PCR reaction included: 7.1 μL molecular grade water, 10 μL 2X Phire Plant PCR Buffer (Thermo Fisher Scientific; includes dNTPs and MgCl_2_), 1 μL of each primer (5 μM), 0.4 μL PHire Hot Start II DNA Polymerase (Thermo Fisher Scientific), and 0.5 μL template DNA. We ran the amplification in a Bio-Rad C1000 Thermal Cycler with the following program: 30s at 98°C, followed by 35-38 cycles (5s at 98°C, 5s at 62°C, 20s at 72°C), 1 min at 72°C. Gel electrophoresis using SYBR Safe (Invitrogen) produced single bands for all products, and no bands in negative controls. We purified PCR products using an Omega Bio-Tek MicroElute® Cycle-Pure Kit and sequenced for both forward and reverse reads (primers ITS5 and ITS4, respectively) on an ABI 3730xl at the University of Illinois Urbana-Champaign Core Sequencing Facility. We could not obtain clean sequencing reads for one morphotype, which we designated as ‘unknown.’

We used Sequencher v5.4 (Gene Codes Corporation) to make base calls, perform quality assessments, and assemble consensus sequences according to 95% ITS sequence similarity, with a minimum of 40% overlap. We performed identification of consensus sequences using BLAST as well as the Ribosomal Database Project (RDP) Bayesian Classifier with both the UNITE and Warcup ITS training sets (Table S2). For BLAST and UNITE classifiers, we reported the top hit. For Warcup, we reported the best named hit, which is provided by default. We assimilated results from across all three classifiers to assign best match taxonomic name for each OTU, and full classifier results and confidence thresholds are reported in Table S2. Sequence data are archived at GenBank under accession numbers XXXXXXXX-XXXXXXXX.

Unexpectedly, we also cultured one bacterial morphotype from soybean leaves, despite using media that selects for the growth of molds and yeasts. It was the most commonly cultured morphotype, isolated more frequently than any fungal morphotype, and found in 58% of colonized tissue fragments and in 70% of leaves that were sampled. To confirm the bacterium could was endophytic to soybean, we inoculated endophyte-free soybean in the growth chamber with the bacterium, and then re-isolated it from leaves of inoculated plants but not endophyte-free control plants (Christian, personal observation).

We extracted total genomic DNA from a single colony using the Qiagen DNeasy Plant Pro kit and incorporated it into a 16S community sequencing run using primers 515F (Parada) and 806R (Apprill). Fluidigm amplification (Brown *et al.*, 2016) and sequencing on the Illumina NovaSeq was performed at the W. M. Keck Center (Urbana, IL, USA). dada2 (v1.14) was used to generate a set of amplicon sequence variants (ASVs) associated with the 209,152 reads generated for this individual sample. 99.944% of those reads were associated with ASVs that were classified as the genus *Methylobacterium* using the SILVA rRNA database (Release 132). All ASVs classified as *Methylobacterium* also clustered together as one OTU at 97% sequence similarity. We thus identified the bacterium as *Methylobacterium* sp., which is consistent with known morphological features of *Methylobacterium* in culture (pink-pigmented colonies). We archived the consensus sequence for this bacterium at GenBank under accession number XXXXXXXXX. *Methylobacterium* is a genus of pink-pigmented facultative methylotrophic bacteria and has generated interest as a fungal pathogen antagonist. However, how *Methylobacterium* affects other non-pathogenic fungi in the plant microbiome is unclear.

### Statistical Analyses – Effects of CO_2_ on soybean plants and endophyte communities

All analyses were performed using R v. 4.0.2 (R Core Team 2020). We used linear mixed effects models to test for the effect of [CO_2_] treatment (elevated vs. ambient) on plant height, LMA, and endophyte abundance (number of isolates per plant), with block included in the model as a random effect (‘lme4’ package, function *lmer*). Significance of random effects was determined using the ‘lmertest’ package (function *rand*). We tested the effect of [CO_2_] on Shannon diversity index using a linear mixed effects model, with LMA included as a fixed effect and block included as a random effect. LMA was included in the model in order to test if LMA predicted endophyte diversity independently from [CO_2_]. The interaction between [CO_2_] treatment and LMA was not significant and not included in the model. Residuals were plotted for all linear mixed models to check for normality. To test the effect of [CO_2_] on endophyte community composition, we used permutational multivariate analysis of variance (PERMANOVA) using distance matrices with the Bray-Curtis dissimilarity index, constraining permutations within block (‘vegan’ package, function *adonis*). We performed the PERMANOVA on all non-singleton OTUs, and Hellinger-transformed the dataset before ordination. Hellinger transformation gives lower weight to species/OTUs with lower counts and many zeros, and is a recommended transformation for species abundance data (Legendre & Gallagher, 2001). To make predictions for how *Methylobacterium* sp. would interact with co-occurring endophytic fungi *in vitro*, we calculated Spearman correlation coefficients using isolate abundance data of *Methylobacterium* sp. and all other OTUs isolated from both ambient and elevated plots (‘spaa’ package, function *sp.pair*).

### Interaction assays

We performed three interaction assay experiments to examine how the presence of *Methylobacterium* sp. affected the growth of soybean-associated fungi *in vitro*. We used the same *Methylobacterium* sp. isolate for all three experiments. It was originally isolated from an ambient [CO_2_] plot.

For Experiment 1, we selected at least one sequenced fungal isolate representing each fungal OTU for the interaction assays, and cultured them on MEA before the start of the experiment. In the instance that we recovered isolates of a single OTU from both the ambient and elevated [CO_2_] plots, we selected one isolate originating from each environment for the interaction assays. In total, we included 31 isolates representing 22 OTUs in the experiment. We also grew an isolate of *Methylobacterium* sp. on MEA, suspended it in water, and adjusted the suspension to an optical density (OD_600_) of 0.1. We plated 15 uL of the bacterial suspension onto 60 mm MEA plates in quadruplicate (protocol modified from Hilber-Bodmer *et al.*, 2017). Twenty-four hours after *Methylobacterium* was plated, we used a sterilized 4mm cork borer to excise fungal plugs from the margin of each fungal plate. We placed a plug in the center of four experimental plates that were pre-inoculated with the *Methylobacterium* (M+), and four control plates that were not inoculated (M−). We incubated plates at room temperature for 10 days or until the margin of at least one replicate fungal culture within a treatment group (on M+ or M− plates) reached the edge of the plate, whichever came first (modified from Hilber-Bodmer *et al.*, 2017). We then photographed plates with a scale and used ImageJ (https://imagej.nih.gov/ij/index.html) to measure the area of each fungal culture.

For Experiment 2, we repeated the methods of Experiment 1, but plated *Methylobacterium* four days before fungal plugs were added.

For Experiment 3, we selected 26 sequenced isolates of the most common fungal OTU, *Colletotrichum* sp. (also the second most common OTU overall, after *Methylobacterium*). 13 isolates originated from plots with elevated [CO_2_] and 13 isolates originated from plots with ambient [CO_2_]. We then conducted interaction assays for these isolates as in Experiment 2 (*Methylobacterium* grew for four days before we added fungal plugs to the plates).

### Statistical analysis – Effects of Methylobacterium on in vitro fungal endophyte growth

All analyses were performed using R v. 4.0.2 (R Core Team 2020). We used ANOVA to test if the presence or absence of *Methylobacterium* affected average colony growth of fungal isolates in all three *in vitro* experiments. Colony area was the response variable, and we included fungal isolate identity (nested within plot type (i.e, ambient vs. elevated [CO_2_]) and *Methylobacterium* presence/absence (and their interaction) as fixed effects. Because the isolate (nested in plot type) by *Methylobacterium* interaction was significant or marginally significant for all experiments (i.e., fungal isolate growth responded to *Methylobacterium* presence but not in the same direction for all isolates), we ran individual t-tests to determine which fungal isolates responded positively, negatively, or did not respond to the presence of *Methylobacterium*. We chose t-tests as planned contrasts instead of a post-hoc comparison (such as a Tukey test) in order to eliminate unnecessary comparisons, and also because the effects of *Methylobacterium* on each fungal isolate were orthogonal. When the average area of fungal growth in the presence of *Methylobacterium* was significantly larger than the average in the absence of *Methylobacterium*, we categorized the *in vitro* interaction as facilitation. When the average area of fungal growth in the presence of *Methylobacterium* was significantly smaller than the average in the absence of *Methylobacterium*, we categorized the *in vitro* interaction as antagonism. In Experiments 1, 2, and 3, there was a high level of variation in growth among different fungal isolates (*P* < 0.001 for all three experiments), as some fungi grow inherently larger or faster in culture compared to others (Table 1).

**Table 1.**
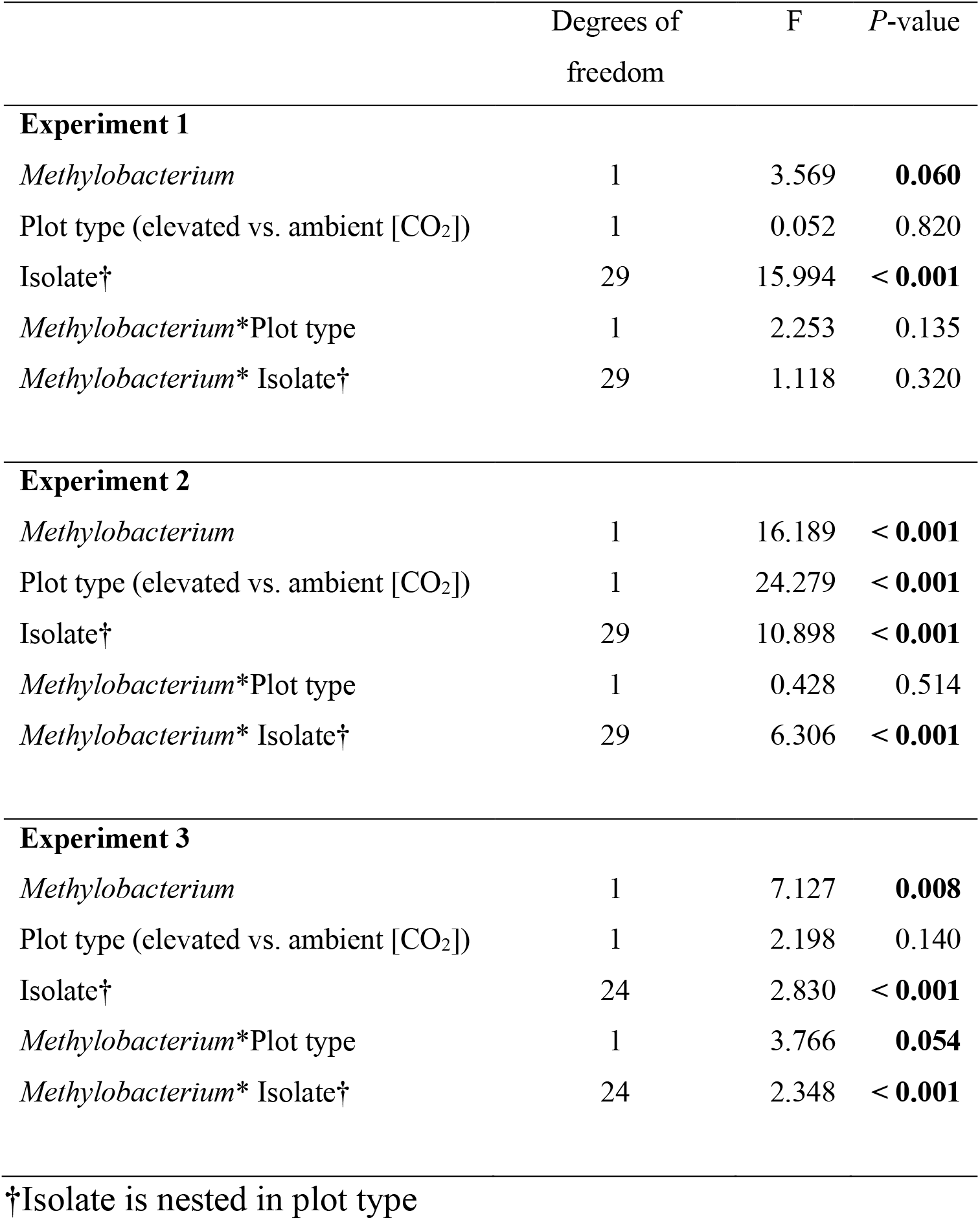
Summary statistics for ANOVAs testing for the effects of *Methylobacterium*, origin of fungal isolation (ambient or elevated [CO_2_] plot type), isolate identity (nested within plot type, and the interaction between *Methylobacterium* and isolate (nested within plot type) on fungal colony area in culture. Statistically significant (p<0.05) and marginally significant (p<0.10) p-values are indicated in bold text. Each fungal isolate was plated with and without *Methylobacterium* in quadruplicate. Experiments 1 and 2 consisted of 31 individual fungal isolates and Experiment 3 consisted of 26 individual fungal isolates.

|  | Degrees of freedom | F | P-value |
| --- | --- | --- | --- |
| **Experiment 1** |  |  |  |
| *Methylobacterium* | 1 | 3.569 | **0.060** |
| Plot type (elevated vs. ambient [CO<sub>2</sub>]) | 1 | 0.052 | 0.820 |
| Isolate† | 29 | 15.994 | **< 0.001** |
| *Methylobacterium* *Plot type | 1 | 2.253 | 0.135 |
| *Methylobacterium* * Isolate† | 29 | 1.118 | 0.320 |
| **Experiment 2** |  |  |  |
| *Methylobacterium* | 1 | 16.189 | **< 0.001** |
| Plot type (elevated vs. ambient [CO<sub>2</sub>]) | 1 | 24.279 | **< 0.001** |
| Isolate† | 29 | 10.898 | **< 0.001** |
| *Methylobacterium* *Plot type | 1 | 0.428 | 0.514 |
| *Methylobacterium* * Isolate† | 29 | 6.306 | **< 0.001** |
| **Experiment 3** |  |  |  |
| *Methylobacterium* | 1 | 7.127 | **0.008** |
| Plot type (elevated vs. ambient [CO<sub>2</sub>]) | 1 | 2.198 | 0.140 |
| Isolate† | 24 | 2.830 | **< 0.001** |
| *Methylobacterium* *Plot type | 1 | 3.766 | **0.054** |
| *Methylobacterium* * Isolate† | 24 | 2.348 | **< 0.001** |

†Isolate is nested in plot type

Finally, to test if results from the *in vitro* experiments predicted endophyte community composition in the field, we regressed the calculated Spearman correlation coefficients between *Methylobacterium* and each fungal isolate in the community survey on the log response ratio (ln(fungal growth in the presence of *Methylobacterium*) – ln(fungal growth in the absence of *Methylobacterium*)) in Experiments 1 and 2.

## Results

### Effects of CO_2_ on soybean plants and endophyte communities

Soybean plants grown under elevated [CO_2_] were taller (df = 1, χ^2^ = 6.052, *P* = 0.014) and had a greater leaf mass per area (LMA) (df = 1, χ^2^ = 7.034, *P* = 0.008) compared to those grown under ambient [CO_2_] (Figure S1). Block (as a random effect) explained a significant amount of variation in height (*P* < 0.001), irrespective of [CO_2_], possibly due to local heterogeneity in environment and soil conditions. Block (as a random effect) did not explain a significant amount of variation in LMA (*P* = 0.198).

Endophyte diversity was positively correlated with LMA (df = 1, χ^2^ = 4.104, *P* = 0.043), a plant trait that was affected by [CO_2_]. However, despite its positive relationship with LMA, endophyte diversity did not differ between plants grown under elevated and ambient [CO_2_] (df = 1, χ^2^ = 1.455, *P* = 0.228). Like diversity, endophyte abundance (i.e., number of isolates per leaf) was not affected by [CO_2_] (df = 1, χ^2^ = 0.005, *P* = 0.945). Endophyte abundance (but not diversity, *P* = 0.167) varied across blocks (*P* = 0.002) (Figure S2), potentially due to variation in plant or site traits across blocks or due to undersampling of endophyte communities (Figure S3), a common constraint in plant microbiome studies (Bullington *et al.*, 2021).

Although endophyte diversity and abundance were overall unaffected by elevated [CO_2_], community composition of foliar endophytic fungi differed between soybean grown under ambient and elevated [CO_2_] (PERMANOVA F = 2.514, R^2^ = 0.029, *P* = 0.043) (Figure 1a, Figure S4). Changes in composition were partially driven by altered abundance of the two most common OTUs under different CO_2_ regimes across blocks (Figure 1b). The best match species names for the most common and second most common OTUs isolated from soybean, respectively, were *Methylobacterium* sp. and *Colletotrichum* sp. 1. Under elevated [CO_2_], the abundance of *Methylobacterium* decreased by 15% compared to under ambient [CO_2_] (Figure 1a,b). In contrast, *Colletotrichum* sp. 1 increased by 58% under elevated [CO_2_] conditions, (Figure 1a,b). Some rare endophytes were lost under elevated [CO_2_] (Figure 1a). There were 8 OTUs isolated under ambient [CO_2_] that were not present under elevated [CO_2_], including a singleton from the genus *Glomerella* (best match name). *Glomerella is* the teleomorph (sexual stage) of *Colletotrichum*, and may be the same genus.

**Figure 1.**
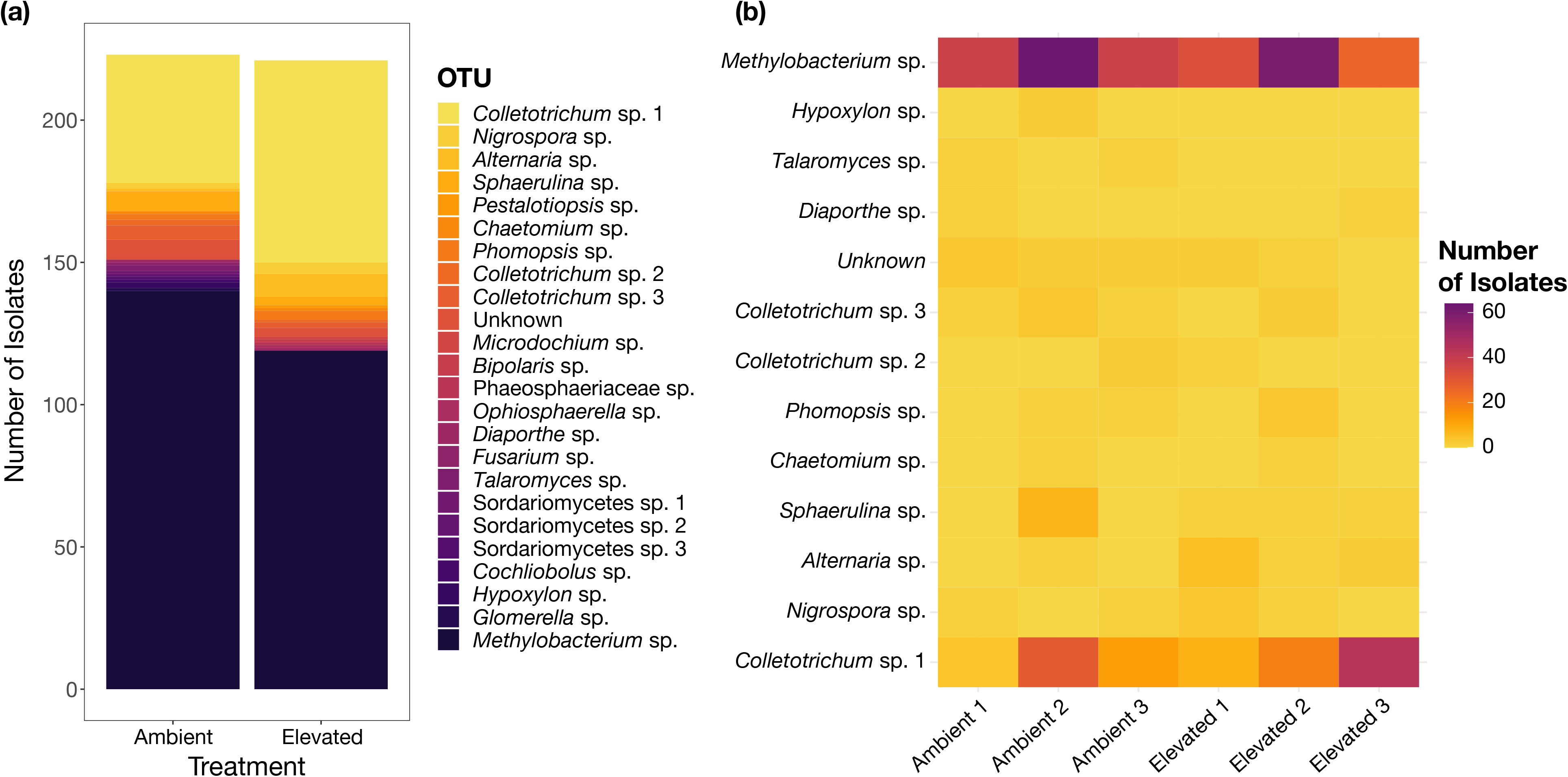
Soybean (*Glycine max*) leaf endophyte communities differed under ambient and elevated [CO_2_] in the field. **a)** The bar plot shows the total abundance distribution of all isolated endophytes labeled with their best match species name. Notable changes under elevated [CO_2_] are a 15% decrease in abundance of *Methylobacterium* sp., the most common endophyte, and a 58% increase in abundance of *Colletotrichum* sp. 1, the second most common endophyte. **b)** The heat map shows abundance (total number of isolates) of each non-singleton isolate distributed across each ambient and elevated plot.

The abundance of *Methylobacterium* was predominantly negatively correlated with other non-pathogenic soybean-associated fungi (Figure 2; for full correlation network see Figure S5). Under ambient [CO_2_], *Methylobacterium* was negatively correlated with 14/18 co-occurring fungi, and under elevated [CO_2_], *Methylobacterium* was negatively correlated with 9/15 fungi (Figure 2; for full correlation network see Figure S5).

**Figure 2.**
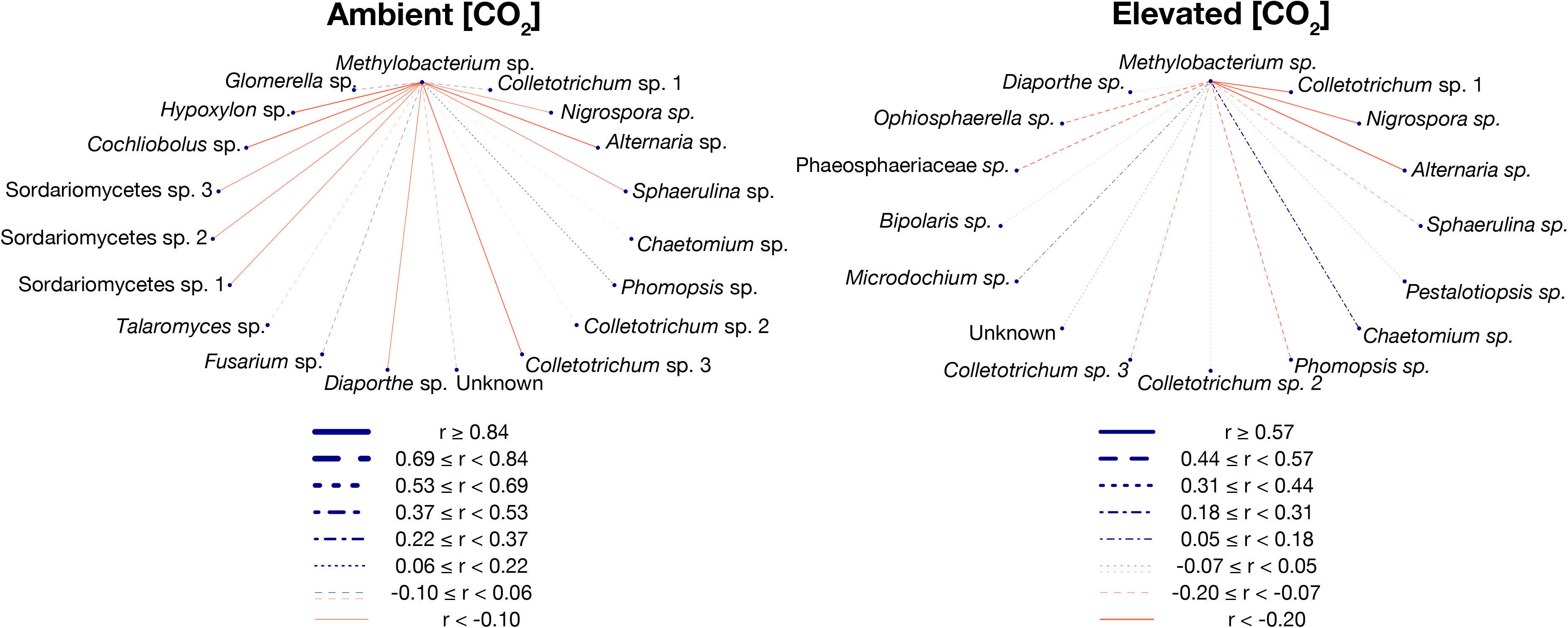
*Methylobacterium* sp., the most commonly isolated endophyte from soybean (*Glycine max*) is predominantly negatively correlated with other co-occurring endophytes. Lines represent Spearman correlation coefficients calculated using absolute abundance of *Methylobacterium* sp. and each OTU within each treatment group. Under ambient [CO_2_], *Methylobacterium* was negatively correlated with 14/18 co-occurring fungi, and under elevated [CO_2_], *Methylobacterium* was negatively correlated with 9/15 fungi. A full network showing all other Spearman correlations between OTUs is found in Figure S5.

### Effects of Methylobacterium on in vitro fungal endophyte growth

In our *in vitro* experiments, we found more significant effects of *Methylobacterium* on fungal growth and larger effect sizes when *Methylobacterium* was more established (Figure 3). When *Methylobacterium* was plated one day before the fungi were added (Experiment 1), the growth of 5 (of 31) fungal isolates was significantly affected by the presence of the *Methylobacterium*, whereas when *Methylobacterium* was allowed to establish for four days (Experiment 2), 12 (of 31) isolates were significantly affected (Figure 3a). Overall, there was a marginally significant effect of *Methylobacterium* on fungal growth in Experiment 1, while the effect in Experiment 2 was highly significant (Table 1; Figure 3). Clearly, more time for *Methylobacterium* to establish in the *in vitro* environment allowed the bacteria to have more pronounced effects, both negative and positive, on fungal growth.

**Figure 3.**
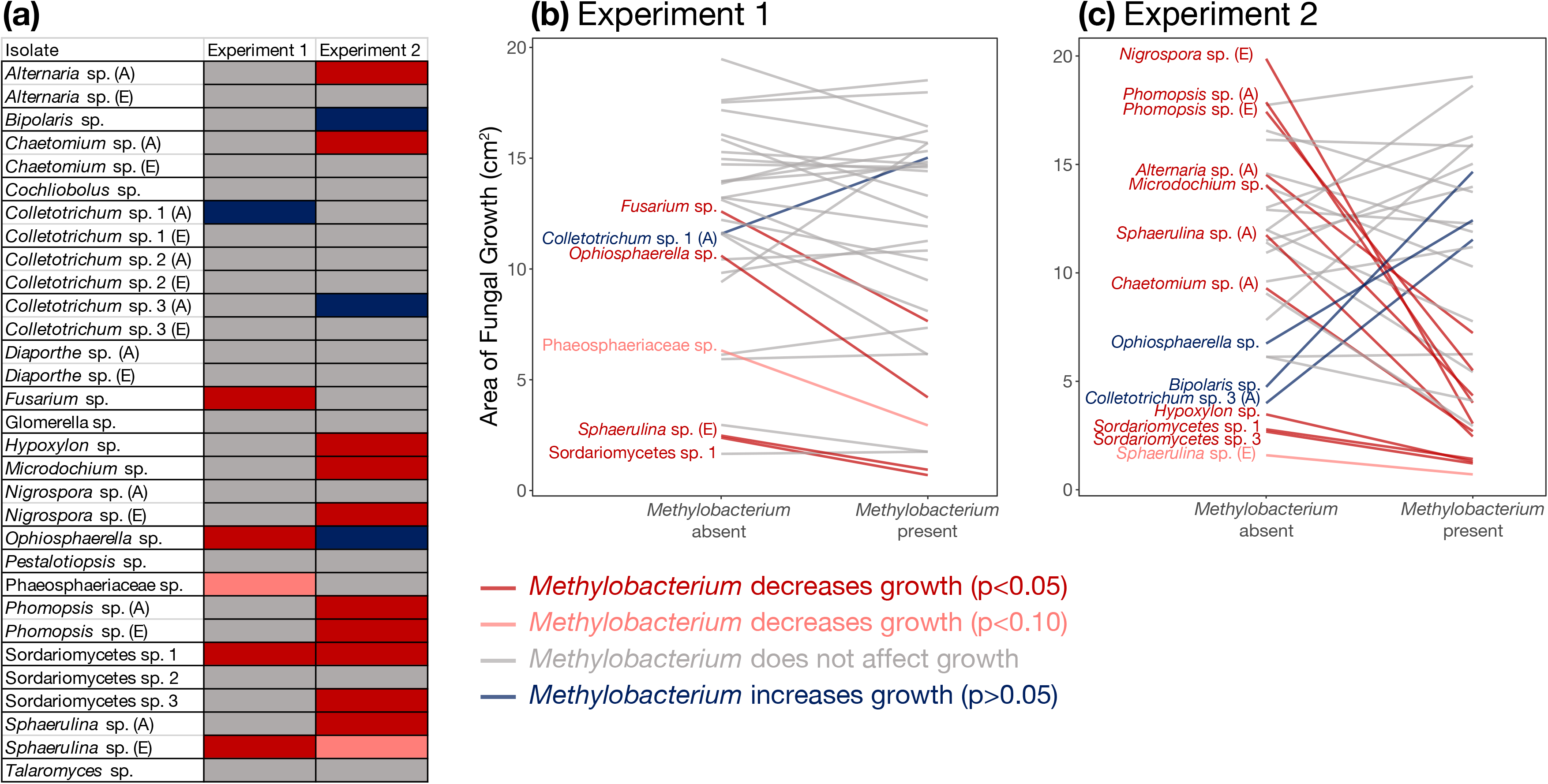
Presence of *Methylobacterium* sp. alters growth of endophytic fungi *in vitro*. 31 fungal isolates representing 22 operational taxonomic units (OTU) were tested. **a)** Table shows Experiment 1, where *Methylobacterium* was plated 1 day before fungal strains were inoculated onto the plate, and Experiment 2, where *Methylobacterium* was plated 4 days in advance. In some cases, two isolates of the same OTU unit were included, one that was originally isolated from an ambient [CO_2_] plot (A) and one that was originally isolated from an elevated [CO_2_] plot (E). Colors represent the direction of the change (increase vs. decrease) and statistical significance of each association assessed using t-tests. **b)** Plot for Experiment 1 shows both the direction and magnitude of change in fungal colony size for each fungal-*Methylobacterium* interaction. Each line represents the average change in colony size of each fungal isolate in the absence and presence of *Methylobacterium.* Colors represent the direction of the change (increase vs. decrease) and statistical significance of each association assessed using t-tests, and isolates that were significant or marginally significant are labeled with best match species names. **c)** Plot for Experiment 2 shows direction and magnitude of change for each fungal-*Methylobacterium* interaction. The number and magnitude of the interactions between *Methylobacterium* and fungi are greater in Experiment 2 compared to Experiment 1.

Of the significant effects of *Methylobacterium* on fungal growth in both Experiments 1 and 2, there were more antagonistic interactions between *Methylobacterium* and fungi than there were interactions in which *Methylobacterium* facilitated fungal growth (Figure 3). Among the five isolates in Experiment 1 that were significantly affected by the presence of *Methylobacterium*, four were negatively affected, while only one fungal isolate was positively affected (Figure 3a, 3b). While most of the significant interactions in Experiment 2 were antagonistic as well (9/12) (Figure 3a, 3c), there were also more instances of bacterial facilitation of fungal growth (Figure 3b, 3c).

The overall preponderance of negative correlations between *Methylobacterium* and co-occurring fungi in the field was supported by the overall dominance of antagonistic interactions in the *in vitro* assays. However, OTU associations *in situ* (i.e., patterns of OTU co-occurrence across samples) did not predict the outcomes of specific interactions between *Methylobacterium* and fungal isolates *in vitro* (Figure 4).

**Figure 4.**
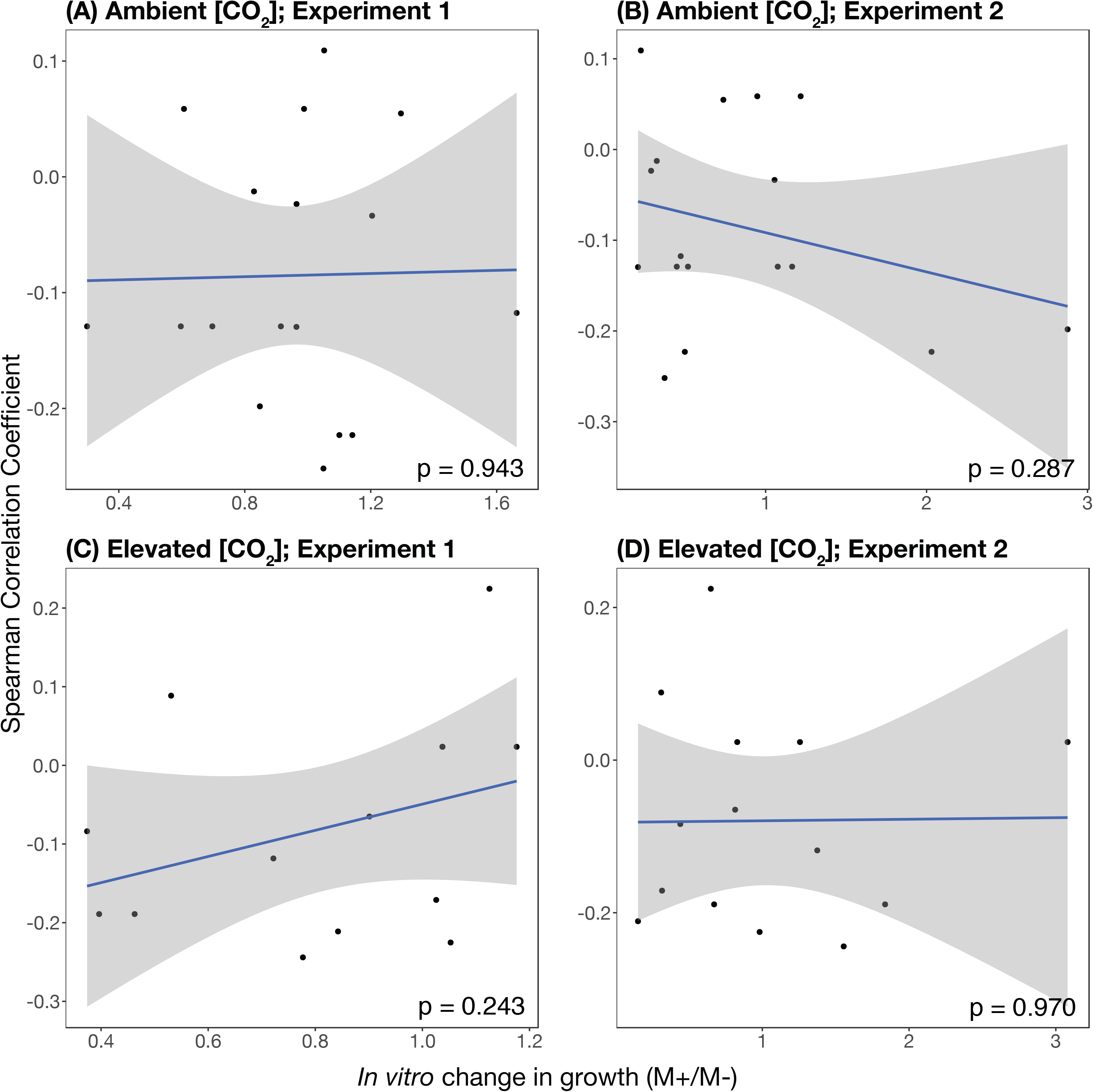
Outcomes of interactions between *Methylobacterium* and fungal isolates *in vitro* did not predict associations between endophytes *in situ* (i.e., patterns of operational taxonomic unit co-occurrence across leaf samples). Spearman correlation coefficients between *Methylobacterium* and OTUs from each [CO_2_] treatment are regressed against the calculated log response ratios (ln(fungal growth in the presence of *Methylobacterium*) – ln(fungal growth in the absence of *Methylobacterium*)). The four different panels represent **(a)** Spearman correlation coefficients from ambient [CO_2_] plots vs. log response ratios of *in vitro* growth in Experiment 1, **(b)** Spearman correlation coefficients from ambient [CO_2_] plots vs. log response ratios of *in vitro* growth in Experiment 2, **(c)** Spearman correlation coefficients from elevated [CO_2_] plots vs. log response ratios of *in vitro* growth in Experiment 1, and **(d)** Spearman correlation coefficients from elevated [CO_2_] plots vs. log response ratios of *in vitro* growth in Experiment 2. Linear regression lines are represented in blue, and the shaded areas represent 95% confidence intervals.

When we tested the response of 26 isolates of *Colletotrichum* sp. 1 to *Methylobacterium* (Experiment 3), we not only found significant variation of growth among those isolates (Table 1) but also that the presence of *Methylobacterium* affected *Colletotrichum* growth (Table 1) by either antagonizing (n = 5) or facilitating (n = 2) growth depending on the isolate (*Methylobacterium* * fungal isolate interaction *P* < 0.001) (Figure 5). Across Experiments 1, 2, and 3, there was as much intra-OTU variation in how fungi responded to *Methylobacterium* as there was inter-OTU variation (Figure 3a, 3b, 5).

**Figure 5.**
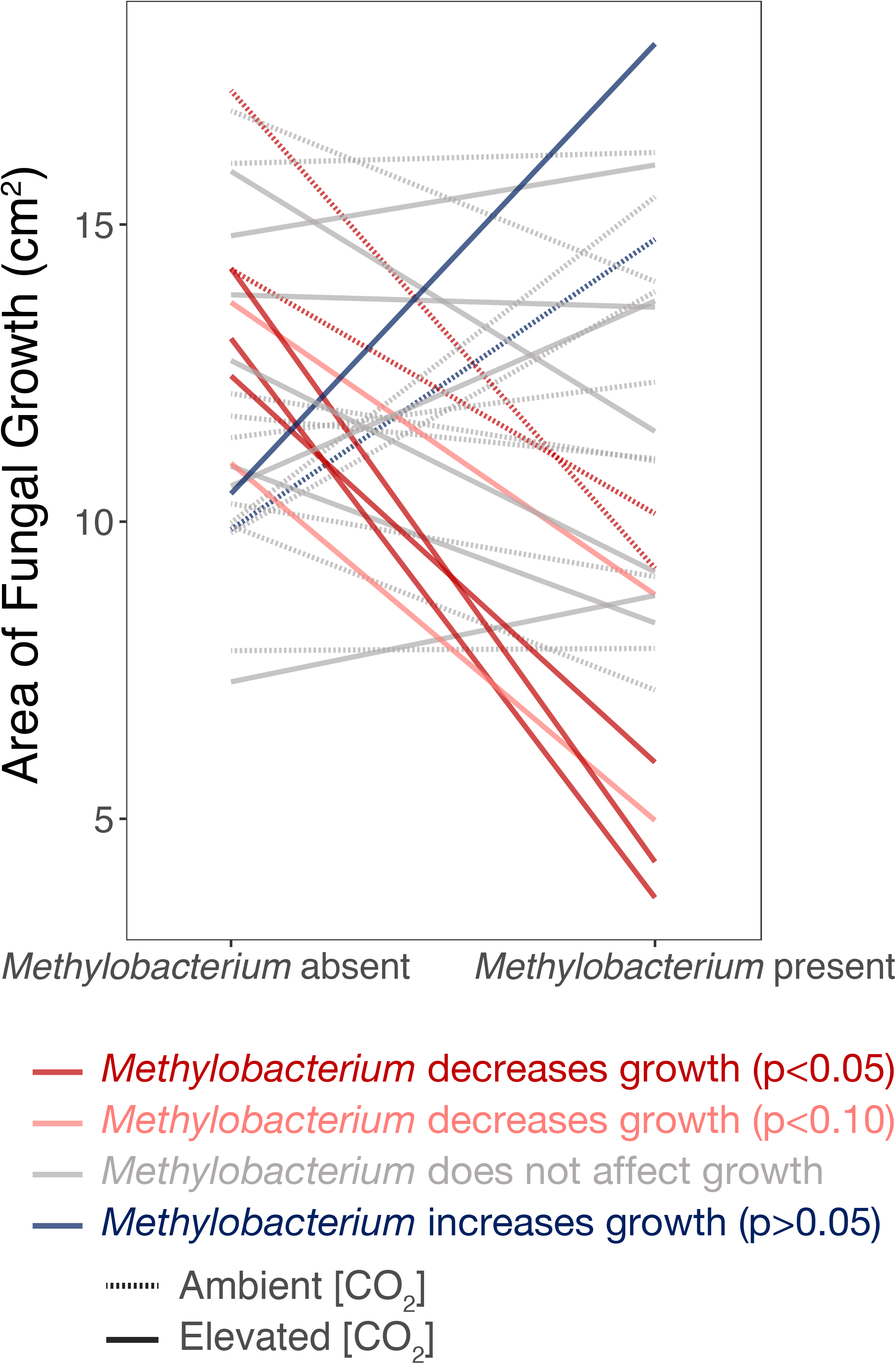
Presence of *Methylobacterium* sp. alters *in vitro* growth of *Colletotrichum* strains (unique isolates clustered as one operational taxonomic unit at 95% similarity) in Experiment 3. Effects were primarily antagonistic but some facilitative interactions were observed. Each line represents the change in growth of various isolates of *Colletotrichum* sp. 1 in the presence and absence of *Methylobacterium.* Color coding of lines represents direction of the change (increase vs. decrease) and statistical significance of each association assessed using t-tests, and line style represents whether isolates originated from an ambient or elevated [CO_2_] plot.

Environmental origin of fungal isolates (isolated from ambient vs. elevated [CO_2_] plots) did not impact *in vitro* growth of fungal isolates in Experiment 1. However, in Experiment 2, fungi that were originally isolated from elevated [CO_2_] plots grew larger in culture than fungi that were originally isolated from plots with ambient [CO_2_], regardless of OTU assignment (Table 1, Figure 6a). *Methylobacterium* had a similar negative effect on fungal growth regardless of plot type of origin (Table 1, Figure 6a). In Experiment 3, the *in vitro* growth of *Colletotrichum* sp. 1 isolates from ambient [CO_2_] plots was not affected by the presence of *Methylobacterium,* whereas there was a marginal negative effect of *Methylobacterium* on the growth of isolates that came from elevated [CO_2_] plots (*Methylobacterium*\*plot type interaction, *P* = 0.054) (Table 1, Figure 6b).

**Figure 6.**
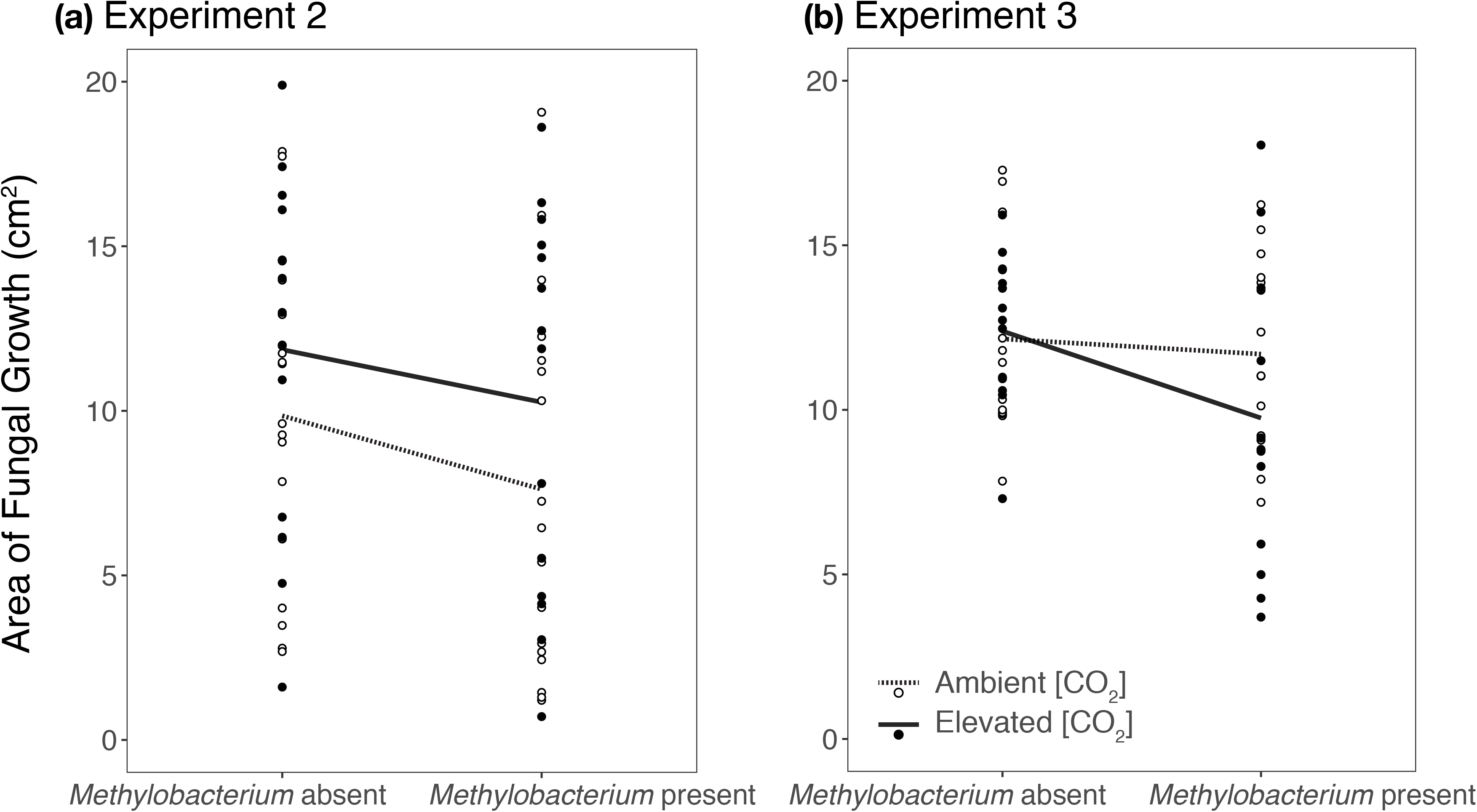
Endophytic fungi isolated from soybean (*Glycine max*) in ambient and elevated [CO_2_] plots behaved differently in culture. **a)** In Experiment 2, taxonomically diverse fungal isolates that came from plots with elevated [CO_2_] grew larger in culture, regardless of *Methylobacterium* sp. presence/absence (*P* < 0.001). **b)** In Experiment 3, growth of *Colletotrichum* sp. 1 strains isolated from ambient [CO_2_] plots was not affected by the presence of *Methylobacterium,* whereas there was a marginal negative effect of *Methylobacterium* on the growth of *Colletotrichum* sp. 1 strains isolated from elevated [CO_2_] plots (*Methylobacterium* * plot interaction, *P* = 0.054).

## Discussion

This study represents the first evidence, to our knowledge, that elevated [CO_2_] affects foliar endophyte community composition in field-grown plants. We found that elevated [CO_2_] reduced the abundance of *Methylobacterium* sp., a common member of the soybean leaf microbiome that was also primarily negatively correlated with other endophytes in the field survey. We then further investigated the ecological impact of *Methylobacterium* by conducting a series of *in vitro* tests that further resolved the positive and negative effects of *Methylobacterium* on co-occurring fungal endophytes. We found that *Methylobacterium* primarily exerted antagonistic effects on co-occurring endophytes *in vitro,* and that intra-OTU variation in response to *Methylobacterium* was comparable to inter-OTU variation. Moreover, we found some evidence that the environment of origin (ambient vs. elevated [CO_2_]) impacted fungal growth and response to *Methylobacterium in vitro.* Here we discuss the implications of our main results for soybean and plants more generally.

While many previous studies have shown how elevated [CO_2_] is changing aspects of plant physiology or ecology, how increasing [CO_2_] affects plant-associated microbial communities is not well understood. We found that soybean plants grown under elevated [CO_2_] were taller and had a greater LMA than those grown under ambient [CO_2_], which is consistent with many previous studies showing that elevated [CO_2_] increases soybean growth responses due to higher carbon assimilation rates (Ainsworth *et al.*, 2002). Endophyte diversity was positively correlated with LMA, potentially suggesting that endophyte community diversity could be influenced by tissue availability, although endophyte diversity was not affected by CO_2_ treatment directly. Previous work across phylogenetically diverse tropical trees suggests the opposite - that endophyte colonization correlates negatively with increasing LMA, since thicker and denser leaf tissue could restrict endophyte hyphal growth (Van Bael *et al.*, 2017). It is possible that there are disparate effects of LMA on endophyte colonization within compared to across host species. It may be that LMA plays a greater role in shaping interspecific differences in endophyte communities compared to intraspecific differences (Tellez *et al.*, 2019).

Elevated [CO_2_] altered the community composition of endophytes in soybean leaves, possibly due to direct effects of [CO_2_] on endophyte colonization or persistence, or alternatively by altering plant traits that selected for a certain suite of endophytes. For example, elevated [CO_2_] decreases stomatal conductance and increases total non-structural carbohydrate concentration (Ainsworth *et al.*, 2002; Ainsworth & Long, 2005). Changes to these traits could impact the colonization and success of certain endophytes over others, as they gain access to the interior of the leaf via stomata (Huang *et al.*, 2018) and consume photosynthesized carbohydrates from host plants (Hardoim *et al.*, 2015). In particular, the abundance of the two most common endophytes in this study (*Methylobacterium* sp. and *Colletotrichum* sp.) was affected by the [CO_2_] treatment.

*Methylobacterium* is a genus of pink-pigmented facultative methylotrophic bacteria that colonizes both the phyllosphere and the rhizosphere of various crops (Dourado *et al.*, 2015; Yoshida *et al.*, 2017), including soybean (Minami *et al.*, 2016), and has generated great interest as an agent for sustainable agriculture (Dourado *et al.*, 2015). *Methylobacterium* and other methylotrophs have been shown to be direct pathogen antagonists (Madhaiyan *et al.*, 2006; Dourado *et al.*, 2015) and can change the overall composition of the endophytic community in ways that reduce disease symptoms (Ardanov *et al.*, 2012). In addition to being a potential biocontrol agent, *Methylobacterium* spp. can also fix nitrogen (Sy *et al.*, 2001) and produce auxins and cytokinins (Ivanova *et al.*, 2001; Meena *et al.*, 2012), ultimately promoting plant nutrient uptake, growth, yield, and land fertility (Kumar *et al.*, 2016). Incorporating *Methylobacterium* and other methylotrophic bacteria into bioinoculants and biofertilizers has been suggested as a way to limit the use of chemical fertilizers, while increasing both plant yield and quality (Kumar *et al.*, 2016). Because of the potential benefits of methylotrophic bacterial colonization, it is of concern that their abundance decreased under elevated [CO_2_] in our study. On the other hand, the abundance of *Colletotrichum* sp. 1 increased substantially under elevated [CO_2_] compared to ambient [CO_2_] (Figure 1). *Colletotrichum* is a diverse genus of fungi, including plant pathogens (Rojas *et al.*, 2010) as well as pathogen antagonists (Arnold *et al.*, 2003; Christian *et al.*, 2019). Although we sampled from healthy, undamaged soybean leaf tissue, it is unclear what functional guild or guilds the *Colletotrichum* in this study belonged to. For example, it could have been a beneficial symbiont, or a latent pathogen that was not yet symptomatic.

The ability to predict microbial interactions in the field from *in vitro* observations, or conversely, to extrapolate ecological relationships using co-occurrence data, is a grand challenge in endophyte ecology and microbiome biology more generally (Whitaker & Bakker, 2019). However, neither of these approaches alone is optimally suited for evaluating endophyte ecology *in situ*. Co-occurrence patterns in the field are shaped not only by interactions between co-occurring species, but also by the consequences of environmental and biotic contexts. Likewise, *in vitro* assays cannot replicate all aspects of the field context in the lab. Combining *in vitro* and *in situ* experiments can help to tease apart the complexity of these systems, even when there are discrepancies between these approaches. For example, Whitaker and Bakker found that *in vitro* interactions between endophytic bacteria and a fungal pathogen did not predict pathogen antagonism *in planta* using a detached wheat head assay (Whitaker & Bakker, 2019). However, endophytic bacteria were antagonistic to the fungal pathogen in culture and also reduced disease severity in an *in vitro* detached spikelet assay, suggesting that some degree of antagonism was conserved in both experimental contexts (Whitaker & Bakker, 2019). Here we present further evidence that while predicting definitive outcomes of pairwise interactions remains elusive, overarching patterns in *in situ* systems can be reflected by *in vitro* assays and vice versa.

Consistent with previous studies that have documented *Methylobacterium* acting as a fungal pathogen antagonist (Dourado *et al.*, 2015), our culture-based survey found that *Methylobacterium* sp. was negatively correlated with most co-occurring fungal endophytes, even though these endophytes were not visibly pathogenic. Negative correlations might represent antagonistic interactions *in situ* such as competition for the same niche, whereas positive correlations between taxa might represent facilitative or mutualistic interactions (Weiss *et al.*, 2016). However, positive or negative associations in a community dataset can be the result of indirect interactions as well, such as shared environmental preferences or other drivers (Weiss *et al.*, 2016). Combining field data with controlled assays can isolate these effects; thus our *in vitro* experiments were designed to resolve the negative and positive effects of *Methylobacterium* on co-occurring fungal endophytes.

In all of our *in vitro* experiments, *Methylobacterium* had predominately inhibitory effects on co-occurring leaf fungi, which is consistent with our *in situ* survey and previous observations that *Methylobacterium* is antagonistic to fungal pathogens of plants (Dourado *et al.*, 2015). Comparing Experiment 1 and Experiment 2, we also found that bacterial establishment time (i.e., how long *Methylobacterium* was allowed to grow in culture before introduction of a fungal colony) affected the outcome of the *Methylobacterium*-fungal interactions. Specifically, we found that *Methylobacterium* reduced the growth of more fungal OTUs, and more strongly, when it was more established. More pronounced antagonistic effects with longer bacterial establishment time could be due to a buildup of antifungal compounds, or simply due to increased resource competition. On the other hand, longer bacterial establishment time could lead to facilitation of fungal growth if the fungi feed directly on bacterial colonies. Previous work has shown that priority effects change the outcome of microbe-microbe interactions *in planta* in ways that impact host disease susceptibility (Adame-Álvarez *et al*., 2014; Leopold & Busby, 2020). While we did not manipulate the order of arrival of the interacting microbes *in vitro*, it is evident that establishment time of early colonizers may be an additional factor that could impact the ecological and functional outcome of microbe-microbe interactions *in planta*. In the development of biocontrol agents and biofertilizers, the fact that *Methylobacterium* could have unintended effects on the composition of the fungal microbiome community (in both positive or negative ways) should be taken into consideration.

Our results from Experiment 3 suggest genetically similar *Colletotrichum* isolates can exhibit significant variation in growth in response to *Methylobacterium* sp. *Colletotrichum* is a functionally diverse genus of fungi, but all of the *Colletotrichum* sp. 1 isolates used in this experiment clustered as the same OTU according to the ITS barcode. The ITS barcode, like the 16S barcode for bacteria, has become the standard marker for illuminating the diversity of microorganisms that inhabit myriad habitats on our planet. However, assessing community composition using standard barcodes may be concealing functional differences among microbial strains.

The phenotypic and functional diversity among strains within a species has long been recognized for microbial symbionts of plants (Parker, 1995; Burdon *et al.*, 1999; Simms *et al.*, 2006; Heath, 2010). Moreover, the importance of incorporating interactions between different levels of biodiversity has been explored in non-microbial communities. In a *Populus* common garden, genetic variation within, rather than between, tree species structured the arthropod community associated with these trees (Whitham *et al.*, 2006). Additionally, genetic diversity in dune plant communities interacted with species diversity to influence overall aboveground biomass production (Crawford & Rudgers, 2012). Intraspecific genetic diversity is expected to have particularly strong effects in communities dominated by one or a few species (Whitham *et al.*, 2006). Not only are endophyte communities often characterized by one or a few very dominant species and many rare species, but individual taxa can also belong to multiple functional guilds beyond that of endophytes, including pathogens and saprotrophs (Zanne *et al.*, 2020). Alternatively, genetically identical endophytes in different physiological stages could have different functional effects on hosts. Whole genome sequencing or multilocus sequence typing paired with more *in vitro* assays would help resolve how intra-OTU genetic diversity quantitatively impacts functional outcomes *in vitro.* Moreover, explicitly assessing if phylogenetic relatedness correlates with functional outcomes *in vitro* could address questions of functional redundancy *in situ*. Uncovering the genetic and functional diversity within plant-associated microbial communities is thus an important direction in microbial ecology, and could be useful in harnessing plant-associated microbiomes for sustainable agriculture in the face of climate change (Busby *et al.*, 2017).

Fungi isolated from elevated vs. ambient [CO_2_] plots differed in colony growth and response to *Methylobacterium*, suggesting that increasing [CO_2_] may affect fungal traits and interactions within the microbiome. In Experiment 2, fungi that were originally isolated from elevated [CO_2_] plots grew larger in culture than fungi that were originally isolated from plots with ambient [CO_2_], regardless of taxonomic identity. It is possible that elevated [CO_2_] selects for fast-growing fungi, as has been seen in the rhizosphere (Dorodnikov *et al*., 2009). In Experiment 3, which tested the response of *Colletotrichum* isolates to *Methylobacterium*, *Methylobacterium* did not impact growth of fungal isolates that originated from ambient conditions, but did have a marginal negative effect on the growth of fungal isolates that came from elevated [CO_2_] plots. If these effects hold in the field, this suggests that *Methylobacterium* could become more antagonistic towards *Colletotrichum* sp. 1 as [CO_2_] increases. Testing for the effects of *Methylobacterium* originally isolated from both ambient and elevated [CO_2_] plots on co-occurring fungal isolates originating from both environments would provide more insight into how increasing [CO_2_] alters interactions between individuals in the soybean leaf microbiome. This is an exciting area of future research connecting plant and microbial ecology, species interactions, and climate change.

In this study, we assessed the differences between endophyte communities of soybean grown under elevated and ambient [CO_2_] using a culture-based methodology and one standard, nutrient-rich growth medium that selects for fungi. It will be useful in future soybean growing seasons to use other types of media as well as culture-independent methodologies to further characterize endophyte communities. Future culture-based approaches could use other types of media to place more emphasis on culturing slow-growing, cryptic fungi, or endophytic bacteria. Illumina sequencing targeting both fungi and bacteria can better quantify how unculturable or rare members of the microbiome respond to elevated [CO_2_] across growing seasons and years, and reveal potential culture biases (e.g., how common *Methylobacterium* sp. is compared to co-occurring bacteria in addition to fungi). Additionally, in our *in vitro* interaction experiments, we quantified how fungi interacted with *Methylobacterium* by measuring fungal growth. Future studies may also consider measuring other fungal traits, including hyphal density, sporulation, or enzymatic production in the presence and absence of *Methylobacterium*. Measuring other fungal traits may lend more insight into how endophytes invest in different life history strategies, and potential trade-offs between growth, resource acquisition, and stress tolerance (Malik *et al*., 2020). Growing endophytes and allowing them to interact with one another in the presence of soybean leaf chemistry would also help refine predictions of how these fungi and bacteria interact *in situ* (Arnold *et al.*, 2003).

Overall, we predict that rising [CO_2_] may change endophyte community composition and potentially function in both crop and non-crop plants, which could also feed back on plant health. Moving forward, we recommend researchers increase efforts to conserve microbial biodiversity in the face of climate change, as cryptic yet important members of communities and ecosystems may be lost. Our *in vitro* experiments highlighted that standard barcodes may obscure functional differences among microbial strains that may have important implications for plants and ecosystems, particularly as the climate continues to change. It will be critical to continue combining field studies with controlled lab experiments to integrate between micro- and field-scale processes. Future research should prioritize experimental inoculations of plants (in controlled settings and the field) with common endophytes to test competitive effects on endophyte communities *in situ* and how changes to endophyte community composition impact host physiology, health, and yield in the face of climate change.

## Acknowledgements

We are grateful to all of the researchers and crew at SoyFACE for setting up and maintaining the experimental plots. Chris Fields and the University of Illinois High Performance Biological Computing facility performed bioinformatics to identify the *Methylobacterium* isolate. Rebecca Batstone provided advice on statistical analyses. Mikus Abolins-Abols assisted with 3-D printing the stencil used to cut leaf fragments. N.C. was supported by a USDA NIFA Postdoctoral Fellowship (AG 2018-67012-27). A.T and B.E.B. were supported by the Phenotypic Plasticity Research Experience for Community College Students (NSF DBI-1559908). X.X. was supported by the Illinois Plant Biology - Fujian Agriculture & Forestry University Study Abroad Program. Additional research support came from the University of Illinois at Urbana-Champaign Campus Research Board (RB18106).

## Author Contribution

N.C., K.D.H., A.T., and B.E.B. designed the experiments. E.A.A. facilitated access to the SoyFACE field site. N.C., B.E.B., A.T., and X.X. conducted the experiments. N.C. analyzed the data and wrote the first draft of the manuscript. K.D.H, P.E.B., and E.A.A. suggested analyses, provided interpretation of the data, and contributed substantial revisions to the manuscript.

## Supporting Information

Additional supporting information may be found in the online version of this article.

**Figure S1.** Soybean (*Glycine max*) height and leaf mass per area differed across plots with ambient and elevated [CO_2_].

**Figure S2.** Effects of block and [CO_2_] treatment on endophyte abundance and diversity.

**Figure S3.** Species accumulation curves across plots, blocks, and [CO_2_] treatments.

**Figure S4.** Endophyte community composition differed between soybean (*Glycine max*) grown under ambient and elevated [CO_2_].

**Figure S5.** Spearman correlation coefficients between operational taxonomic units isolated from soybean (*Glycine max*) grown under ambient and elevated [CO_2_].

**Table S1** Meteorological data for the 2018 SoyFACE growing season

**Table S2** Best match taxonomic assignments of each fungal operational taxonomic unit

**Table S3** Endophyte community data

**Table S4** *In vitro* interaction assay data: Experiments 1-2

**Table S5** *In vitro* interaction assay data: Experiment 3

