## Supplemental Material for "Elevated CO_2_ reduces a common soybean leaf endophyte"

**Table S2** Best match taxonomic assignments of fungal operational taxonomic unit using the Ribosomal Database Project (RDP) Bayesian Classifier with both the UNITE and Warcup ITS training sets, as well as NCBI Nucleotide BLAST.

| Fungal  OTU | UNITE | Warcup | BLAST |
| --- | --- | --- | --- |
| 1 | Glomerella_tucumanensis\|SH229539.06FU [100%] | Colletotrichum spaethianum[70%] | Colletotrichum spaethianum [100.00%] |
| 2 | Trichosphaeriales_sp\|SH237073.06FU [100%] | Nigrospora oryzae[98%] | Nigrospora sp. [99.82%] |
| 3 | Alternaria_eichhorniae\|SH224789.06FU [100%] | Alternaria alternata[58%] | Alternaria tenuissima [100.00%] |
| 4 | Sphaerulina_pseudovirgaureae\|SH212655.06FU [100%] | Mycosphaerella rubi[84%] | Septoria sp. [100.00%] |
| 5 | Pestalotiopsis_rhododendri\|SH210426.06FU [100%] | Pestalotiopsis olivacea[78%] | Pestalotiopsis neglecta [99.00%] |
| 6 | Ascomycota_sp\|SH234918.06FU [99%] | Chaetomium coarctatum[71%] | Chaetomium sp. [98.42%] |
| 7 | Diaporthales_sp\|SH194734.06FU [100%] | Phomopsis longicolla [100%] | Diaporthe longicolla [99.47%] |
| 8 | Glomerellaceae_sp\|SH233480.06FU [100%] | Glomerella magna [62%] | Colletotrichum gloeosporioides [99.82%] |
| 9 | Colletotrichum_chlorophyti\|SH229546.06FU [100%] | Colletotrichum chlorophyte [100%] | Colletotrichum chlorophyti [100.00%] |
| 10 | Microdochium_sp_E9023a\|SH216936.06FU [100%] | Microdochium bolleyi [93%] | Microdochium sp. [97.57%] |
| 11 | Bipolaris_microstegii\|SH224794.06FU [97%] | Cochliobolus sativus [56%] | Bipolaris sorokiniana [97.93%] |
| 12 | Chaetosphaeronema_sp\|SH231242.06FU [26%] | Phaeosphaeria phragmitis [63%] | Ophiosimulans sp. [95.94%] |
| 13 | Fungi_sp\|SH192549.06FU [94%] | Ophiosphaerella agrostis [100%] | Uncultured fungus clone [98.22%] |
| 14 | Diaporthe_caulivora\|SH194728.06FU [100%] | Diaporthe caulivora [100%] | Diaporthe caulivora [100.00%] |
| 15 | Ascomycota_sp\|SH219457.06FU [95%] | Fusarium avenaceum [59%] | Fusarium reticulatum [98.19%] |
| 16 | Trichocomaceae_sp\|SH225848.06FU [100%] | Talaromyces trachyspermus [100%] | Talaromyces trachyspermus [98.38%] |
| 17 | Sordariomycetes_sp\|SH212494.06FU [89%] | Colletotrichum dracaenophilum [20%] | Uncultured endophytic fungus clone [94.42%] |
| 18 | Arthrinium_pterospermum\|SH233390.06FU [40%] | Discosia sp 2 KT_2010 [12%] | Fungal endophyte strain [97.85%] |
| 19 | Sordariomycetes_sp\|SH212485.06FU [99%] | Podospora fimiseda [7%] | Fungal sp. strain [97.98%] |
| 20 | Pleosporaceae_sp\|SH224808.06FU [100%] | Cochliobolus pallescens [79%] | Uncultured fungus clone [98.76%] |
| 21 | Hypoxylon_submonticulosum\|SH236019.06FU [100%] | Hypoxylon monticulosum [60%] | Hypoxylon submonticulosum [98.61%] |
| 22 | Glomerella_graminicola\|SH229544.06FU [100%] | Colletotrichum navitas [36%] | Glomerella graminicola [99.65%] |

Percentages given in brackets indicate confidence threshold (for UNITE and Warcup) and percent identify (for BLAST) compared to best match.

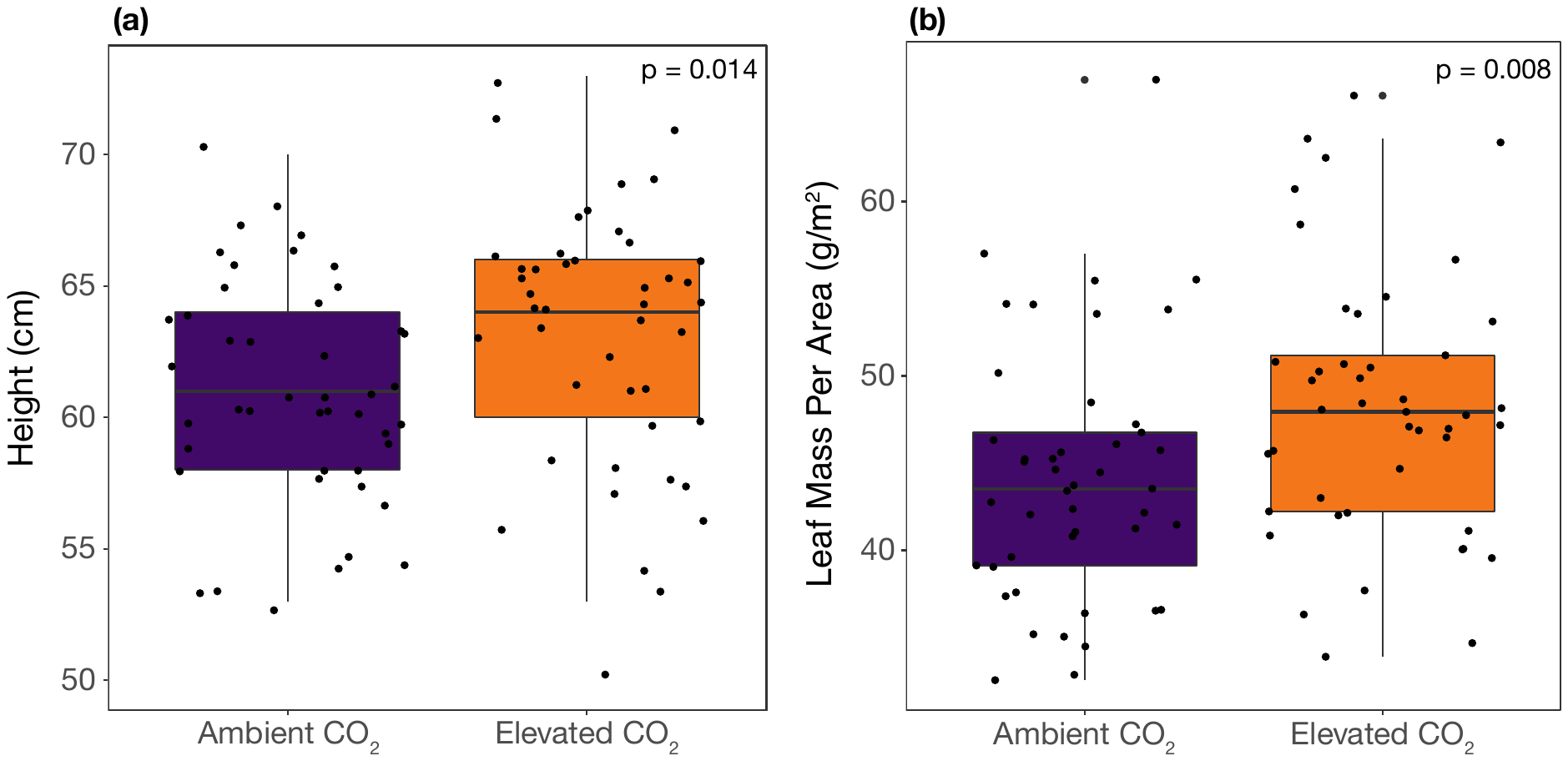

**Fig. S1** Soybean (*Glycine max*) (a) height and (b) leaf mass per area differed across plots with ambient and elevated [CO_2_]. Boxes show first and third quartiles with the median as a heavy line, and whiskers extend to 1.5 times the interquartile range.

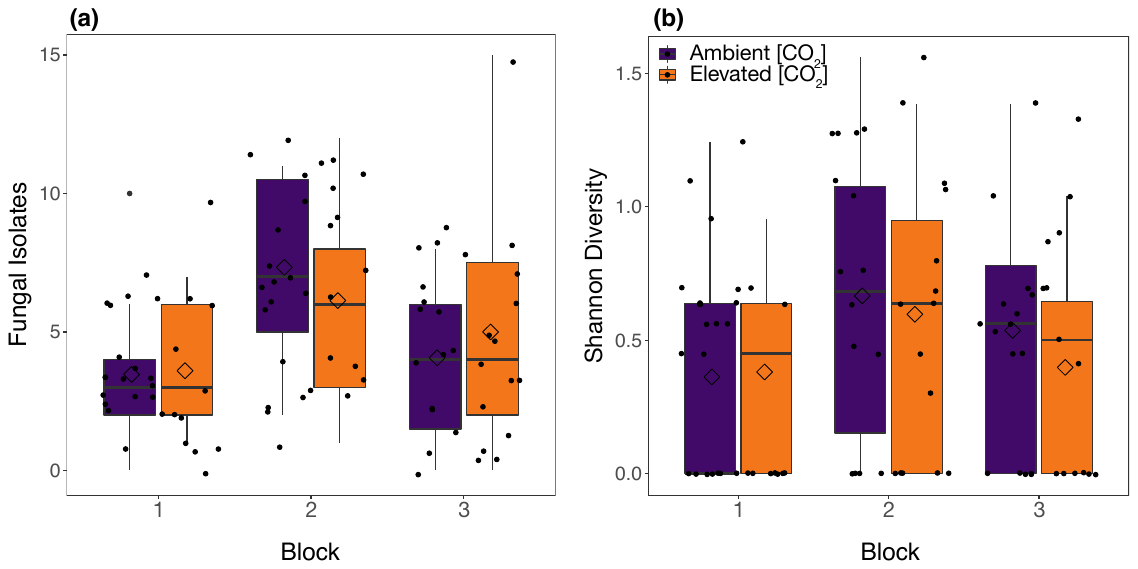

**Fig. S2** Effects of block and [CO_2_] treatment on endophyte abundance and diversity. a) Number of endophyte isolates from soybean (*Glycine max*) was not affected by [CO_2_] (*P* = 0.945) but differed across paired field plots (block as random effect, *P* = 0.002). b) Endophyte diversity was not affected by [CO_2_] (*P* = 0.228) nor differed across blocks (*P* = 0.167). Boxes show first and third quartiles with the median as a heavy line and mean as diamond, and whiskers extend to 1.5 times the interquartile range.

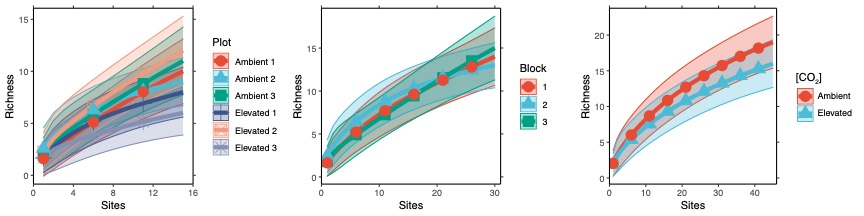

**Fig. S3** Species accumulation curves across plots, blocks, and [CO_2_] treatments.

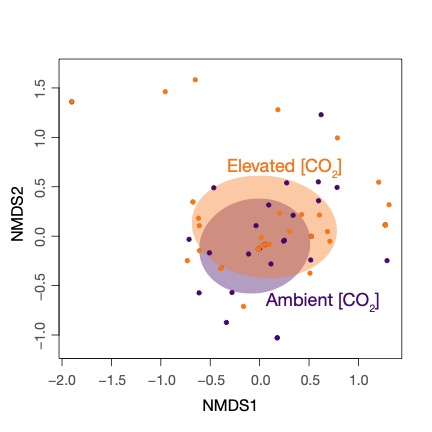

**Fig. S4** Endophyte community composition differed slightly between soybean (*Glycine max*) grown under ambient and elevated [CO_2_] (R^2^ = 0.029, *P* = 0.043).

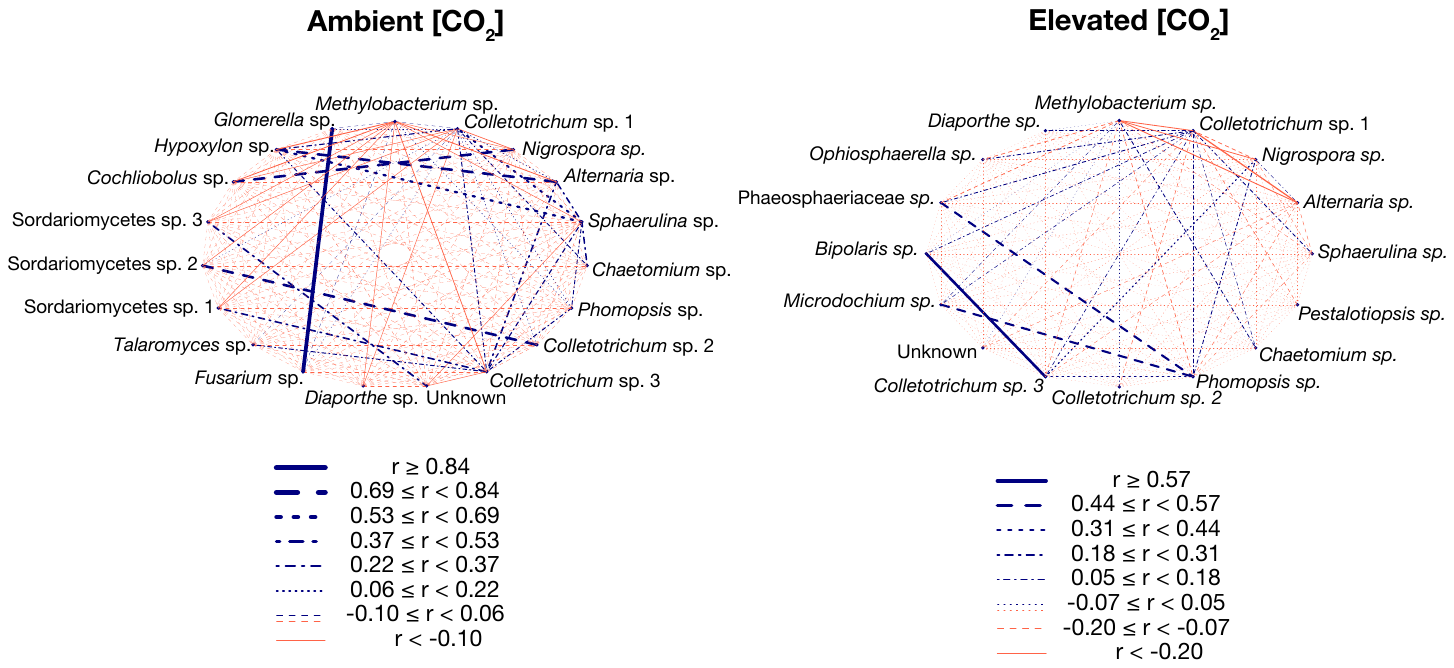

**Fig. S5** Spearman correlation coefficients between operational taxonomic units (OTU) isolated from soybean (*Glycine max*) grown under ambient and elevated [CO_2_]. Lines represent Spearman correlation coefficients calculated using absolute abundance of non-singleton OTUs from each plot type (elevated vs. ambient [CO_2_].
